# Phylogenomics Sheds Light on the population structure of *Mycobacterium bovis* from a Multi-Host Tuberculosis System

**DOI:** 10.1101/2021.04.26.441523

**Authors:** Ana C. Reis, Liliana C.M. Salvador, Suelee Robbe-Austerman, Rogério Tenreiro, Ana Botelho, Teresa Albuquerque, Mónica V. Cunha

## Abstract

Molecular analyses of *Mycobacterium bovis* based on spoligotyping and Variable Number Tandem Repeat (MIRU-VNTR) brought insights into the epidemiology of animal tuberculosis (TB) in Portugal, showing high genotypic diversity of circulating strains that mostly cluster within the European 2 clonal complex. The genetic relatedness of *M. bovis* isolates from cattle and wildlife have also suggested sustained transmission within this multi-host system. However, while previous surveillance highlighted prevalent genotypes in areas where livestock and wild ungulates are sympatric and provided valuable information on the prevalence and spatial occurrence of TB, links at the wildlife-livestock interfaces were established mainly via genotype associations. Therefore, evidence at a local fine scale of transmission events linking wildlife hosts and cattle remains lacking. Here, we explore the advantages of whole genome sequencing (WGS) applied to cattle, red deer and wild boar isolates to reconstruct the evolutionary dynamics of *M. bovis* and to identify putative pathogen transmission events. Whole genome sequences of 44 representative *M. bovis* isolates, obtained between 2003 and 2015 from three TB hotspots, were compared through single nucleotide polymorphism (SNP) variant calling analyses. Consistent with previous results combining classical genotyping with Bayesian population admixture modelling, SNP-based phylogenies support the branching of this *M. bovis* population into five genetic clades, three with geographic specificities, as well as the establishment of a SNPs catalogue specific to each clade, which may be explored in the future as phylogenetic markers. The core genome alignment of SNPs was integrated within a spatiotemporal metadata framework to reconstruct transmission networks, which together with inferred secondary cases, further structured this *M. bovis* population by host species and geographic location.

WGS of *M. bovis* isolates from Portugal is reported for the first time, refining the spatiotemporal context of transmission events and providing further support to the key role of red deer and wild boar on the persistence of animal TB in this Iberian multi-host system.

## 1. Introduction

*Mycobacterium bovis* is an important pathogen, responsible for causing animal tuberculosis (TB) in livestock and wildlife vertebrates, as well as in humans (Brites et al., 2018; Gagneux, 2018). Cattle (*Bos taurus*) is the main livestock affected species, while several reports evidence the importance of the livestock-wildlife interface for disease maintenance (Corner, Murphy, & Gormley, 2011; Fitzgerald & Kaneene, 2012; M. V. Palmer, 2007; Mitchell V. Palmer, Thacker, Waters, Gortázar, & Corner, 2012). In the Iberian Peninsula, red deer (*Cervus elephus*) and wild boar (*Sus scrofa*) have both been implicated in the transmission of *M. bovis* to cattle via direct and indirect routes and in pathogen persistence across ecosystems, depending on the specificities of the epidemiological scenario and the ecological relationships established by the hosts (Barasona, Torres, Aznar, Gortázar, & Vicente, 2017; Cunha et al., 2012; Gortázar et al., 2008; Naranjo, Gortázar, Vicentea, & de la Fuente, 2008; Santos et al., 2009; Vieira-Pinto et al., 2011). The presence of maintenance hosts in the wild is associated with difficulties in the success of test and slaughter schemes implemented in the cattle population, but it also brings concerns regarding wildlife welfare, biodiversity and public health.

To date, works with reference to *M. bovis* molecular characterization in Portugal have been based on the analysis of repetitive genomic regions, namely spoligotyping (spacer oligonucleotide typing) and MIRU-VNTR (*Mycobacterial Interspersed Repetitive Units-Variable Number Tandem Repeats*). The focus has been placed over isolates from cattle, red deer and wild boar from TB hotspot areas located in the central and south regions of the country (Cunha et al., 2012; Duarte, Domingos, Amado, Cunha, & Botelho, 2010; Reis, Tenreiro, Albuquerque, Botelho, & Cunha, 2020). These works provided evidences for *M. bovis* population diversity and structure, highlighting the main genotypes across host species and regions, as well as intra- and inter-specific transmission (Cunha et al., 2012; Duarte et al., 2010; Reis et al., 2020). However, these molecular typing approaches have explored epidemiological links via genotype associations, which are not sufficiently discriminatory to accurately assess transmission at a fine-scale, nor to gain insights on the roles exerted by different species in a multi-host system. But understanding the evolutionary processes driving transmission among sympatric wildlife reservoirs and livestock populations is crucial for an effective management of animal TB in an endemic system.

The progressive application of whole genome sequencing (WGS) to infectious disease systems has resulted in unprecedented advances in the ability to resolve epidemiological information at different scales. WGS data provides higher discriminatory power than classical molecular approaches for resolving complex outbreak situations, allowing a finer definition of the spatiotemporal context in which pathogen spread and persistence occurs. WGS also aids in the identification of the infection source, the establishment of epidemiological links, and the reconstruction of transmission chains (Biek et al., 2012; Glaser et al., 2016; Price-Carter et al., 2018).

When considering the livestock-wildlife interface, WGS has been used to demonstrate the close genetic relationship among *M. bovis* isolates recovered from sympatric cattle and wildlife populations, in different epidemiological settings, including UK (Biek et al., 2012; Crispell et al., 2019; Trewby et al., 2016), Ireland (Crispell et al., 2020), New Zealand (Crispell et al., 2017; Price-Carter et al., 2018) and United States of America (Glaser et al., 2016; Salvador et al., 2019). In this context, single nucleotide polymorphisms (SNPs) emerged as good phylogenetic markers, helping in the definition of *M. bovis* population structure, having been recently used to define four *M. bovis* lineages, and to inform transmission models (Biek et al., 2012; J. Guerra-Assunção et al., 2015; Price-Carter et al., 2018). When placed together with data concerning the time needed for this slow growing bacterium to accumulate new SNPs, this information can provide temporal clues on the emergence and divergence of specific genotypes.

With the aim to improve knowledge on *M. bovis* population structure and transmission within and across TB hotspots in Portugal, WGS of 44 *M. bovis* obtained between 2003 and 2015 from cattle, red deer and wild boar was completed. These isolates were selected as being representative of the *M. bovis* population diversity in those areas, which was previously assessed by a large-scale genotyping study that combined standard genotyping techniques (spoligotyping and MIRU-VNTR) with Bayesian clustering (Reis et al., 2020). The methodological framework aimed to: (1) identify phylogenetic clades and build a catalogue of SNPs that may be used as specific molecular markers of each clade; (2) explore how specific nucleotide differences are associated with distinct host and/or geographic regions; (3) disclose transmission networks for global and local datasets from Portugal; and (4) infer infectivity in distinct host species and geographic locations.

## 2. Methods

### 2.1. *M. bovis* isolates dataset

The 44 *M. bovis* isolates used in this work were recovered from cattle (*n*=16), red deer (*n*=16) and wild boar (*n*=12) from TB hotspot areas in Portugal that encompass the districts (administrative level sample unit) of Castelo Branco, Portalegre and Beja, which are located in inner central and south of the mainland territory (Supplementary Fig. 1 and Supplementary Table 1). This dataset was selected for WGS from a wider *M. bovis* dataset recovered in Portugal (*n*=487) that was previously submitted to molecular characterization by classical genotyping techniques, namely spoligotyping and 8 *loci* MIRU-VNTR (Supplementary Fig. 1) (Reis et al., 2020). To represent the population genetic diversity, two selection criteria were applied: first, isolates should represent major spoligotyping-MIRU type groups and adjacent variants; and second, they should cover different host species, geographic regions and temporal/epidemiological contexts (using year of *M. bovis* isolation as proxy) (Supplementary Fig. 1).

### 2.2. Ethical Approval

The *M. bovis* dataset analysed here was selected for WGS from a wider *M. bovis* dataset recovered in Portugal (Reis et al. 2020) in the scope of official control plans for animal TB. No animals were sacrificed for the purposes of this study. Isolates were obtained in the national reference laboratory of animal tuberculosis (INIAV, IP) from animal samples either presenting TB-compatible lesions during official inspection and/or animal samples from reactor cattle submitted to official standard screening test for TB [the single intradermal comparative cervical tuberculin (SICCT) test]. None of the authors were responsible for the death of any animals nor were any samples used in the study collected by the authors. All applicable institutional and/or national/international guidelines for the care and use of animals have been followed.

### 2.3. DNA extraction

Bacteriological culture was performed as described by Reis et al. (2020). Frozen culture stocks of 44 *M. bovis* were successfully re-cultured on Middlebrook 7H9 (Difco) medium supplemented with 5% sodium pyruvate and 10% ADS enrichment (50 g albumin, 20 g glucose, 8.5 g sodium chloride in 1 L water), at 37 °C, on a level 3 biosecurity facility.

After four weeks’ growth, the culture medium was renewed and the cultures were monitored regularly until growth was observed. Cells were harvested, centrifuged and the culture pellet was re-suspended in 500 μL PBS and inactivated by heating at 99°C for 30 minutes. After centrifugation, the supernatants were stored at −20°C until WGS.

### 2.4. Whole-genome sequencing and SNP analysis

The genomic DNA was sequenced using the Illumina Genome Analyser, according to the manufacturer’s specifications with the paired-end module attachment. Forty-two samples were sequenced by MiSeq technology (2×250 pb) at the United States Department of Agriculture (USDA, USA) and the remaining two by HiSeq (2×150 pb) (Eurofins, Germany).

The vSNP pipeline, currently available at https://github.com/USDA-VS/vSNP, was used to process the FASTQ files obtained from Illumina sequencing. Briefly, reads were aligned to the *M. bovis* AF2122/97 reference genome (NCBI accession number NC_002945.4), using BWA and Samtools (Li & Durbin, 2009; Li et al., 2009). Base quality score recalibration, SNP and indel (insertion or deletion) discovery were applied across all isolates using standard filtering parameters or variant quality score recalibration according to Genome Analysis Toolkit (GATK)’s Best Practices recommendations (Depristo et al., 2011; Mckenna et al., 2010; Van der Auwera et al., 2014). Results were filtered using a minimum SAMtools quality score of 150 and AC = 2.

Integrated Genomics Viewer (IGV) (version 2.4.19) (Thorvaldsdóttir, Robinson, & Mesirov, 2012) was used to visually validate SNPs and positions with mapping issues or alignment problems. SNPs that fell within Proline-Glutamate (PE) and Proline-Proline Glutamate (PPE) genes were filtered from the analysis, as well as indels.

The raw data is deposited in a public domain server at the National Centre for Biotechnology Information (NCBI) SRA database, under BioProject accession number PRJNA682618.

### 2.5. Phylogenetic analysis

Validated and polymorphic SNPs were concatenated, resulting into a single 1842-nt sequence. MEGA (Molecular Evolutionary Genetics Analysis, version 7.0) (Kumar, Stecher, & Tamura, 2016) was used to conduct phylogenetic analysis, using the maximum likelihood method with 1,000 bootstrap inferences.

Distribution of pairwise SNP distance were obtained by applying the Hamming distance, using the library *ape*, and the corresponding heatmap was obtained by library *gplots*, both in R statistical package (Paradis, Claude, & Strimmer, 2004).

### 2.6. Transmission mapping

The *SeqTrack* library in R statistical package (Jombart, Eggo, Dodd, & Balloux, 2011) was used to infer total and local transmission networks. This library builds a network, minimizing the genomic distance between links and keeping the disease onset dates coherent, reconstructing the most plausible genealogy for a determined group of isolates. Matrices with information concerning host species and geographic region were added to determine the transmission pathway (Jombart et al., 2011).

The geographic coordinates of *M. bovis* isolates (*n*=36, for which georeferenced information was complete) allowed the definition of two hotspot areas that were further analysed in local networks (Supplementary Fig. 2). Each *M. bovis* isolate from cattle was assigned to the geographic coordinates of the corresponding livestock herd; isolates from hunter-harvested red deer and wild boar were associated to the centroids of officially-delimited hunting areas, and no substantial change in geographic coordinates was assumed over the study period. Therefore, a total network with all *M. bovis* and two local networks, one for Castelo Branco district (*n*=16) and another one for Portalegre (*n*=12) were performed.

Putative recent transmission events were established based on a maximum inter-isolate pairwise SNP distance of three SNPs within 5-year range (Crispell et al., 2020).

The number of secondary cases was inferred, based on the number of established links between isolates and thereafter grouped by host species and geographic region, being displayed in a violin density plot. The statistical significance between the inferred number of secondary cases and host species or geographic region were tested using a one-way ANOVA test. Results were considered to be significant if p < 0.05. All the statistical tests were performed in R software environment (version 3.4.6).

### 2.7. Phylogeographic analysis

A phylogeographic analysis was performed with a combined approach including Gephi version 0.9.2 (Bastian & Heymann, 2009) and QGIS (Quantum GIS development Team 2018, version 3.10).

A network analysis to explore the relationships established between *M. bovis* isolates, using as node each *M. bovis* and as connection lines the number of shared SNPs, was performed in Gephi for four cumulative temporal periods, and plotted in a map with QGIS. Four networks were generated with a progressive temporal interval of three years between each other, resulting in the definition of period 1 (2003-2006), period 2 (2003-2009), period 3 (2003-2012) and period 4 (2003-2015). These temporal windows follow the variation of animal TB epidemiological indicators in Portugal, considering herd prevalence values in cattle and the surveillance measures implemented in wildlife (Supplementary Fig. 3). Thus, in periods 1 and 2, the values of herd prevalence steadily decreased; in period 3, an increase in herd prevalence was registered and carcass examination of hunted big game species became mandatory in the epidemiological risk area that is under analysis in this study; and, finally, in the last years added to the timeline, a decrease in herd prevalence was observed (Supplementary Fig. 3). The connection lines were established based on absolute values of SNPs; no statistical transformation was performed.

A total of 36 *M. bovis* had information concerning geographic coordinates of the corresponding livestock herd or officially-delimited hunting area. For eight *M. bovis* strains [cattle (*n*=5), wild boar (*n*=2), red deer (*n*=1)], the specific geographic coordinates were absent, so the coordinates were assumed as the centroid of the geographic region, so that all *M. bovis* could be plotted in the map. No substantial change in geographic coordinates was assumed over the study period.

### 2.8. Isolation by distance analysis

A Mantel test was conducted in the R software environment package *ade4* (Dray & Dufour, 2007) to assess correlation between spatial and SNP distances, using 10,000 permutations to assess significance. Only the isolates with geographical coordinates were included in this analysis. The Mantel test was applied to the total dataset (*n*=36) and to local networks of Castelo Branco (*n*=16) and Portalegre (*n*=12).

### 2.9. Molecular dating

The temporal structure of the sequences was explored with *Tempest* (TEMPoral Exploration of Sequences and Trees, version 1.5.1) (Rambaut, Lam, Carvalho, & Pybus, 2016) and with a date randomization test (DRT) with LSD (Least-Squares Dating, version 0.3-beta) (To, Jung, Lycett, & Gascuel, 2015). In the approach with *Tempest*, a phylogenetic tree performed with the *M. bovis* AF2122/97 reference genome as outgroup was used as input.

The least square method implemented in LSD v0.3-beta (To et al., 2015) was applied to estimate the molecular clock rate in the observed data and to perform a DRT with 20 randomized datasets. The QPD (quadratic programming dating) algorithm was used, allowing to estimate the position of the root (option -r as) and calculating the confidence intervals (options -f 100 and -s) (To et al., 2015).

## 3. Results

### 3.1. SNP-based genotyping and phylogenetic analyses

The sequence reads of 44 *M. bovis* whole genomes representing the genetic diversity of strains circulating in TB hotspots in Portugal were mapped to the assembled reference genome of *M. bovis* AF2122/97 (NC_002945.4) (Supplementary Table 1). The average depth of coverage and genome coverage was 93.6 and 99.57%, respectively (Supplementary Table 2). The SNP alignment had a total of 1842 polymorphic positions, being the majority (86.5%) located in coding regions.

The phylogenetic distribution of SNPs grouped *M. bovis* into five related clades, each one with more than 100 clade-defining SNP sites, i.e. polymorphic positions specifically found within each clade member (Fig. 1a and Table 1). Clades were named from A to E, being clade A the largest, counting with 14 *M. bovis* genomes, while clade C is the smallest, encompassing only three (Fig. 1a).

**Figure 1.**
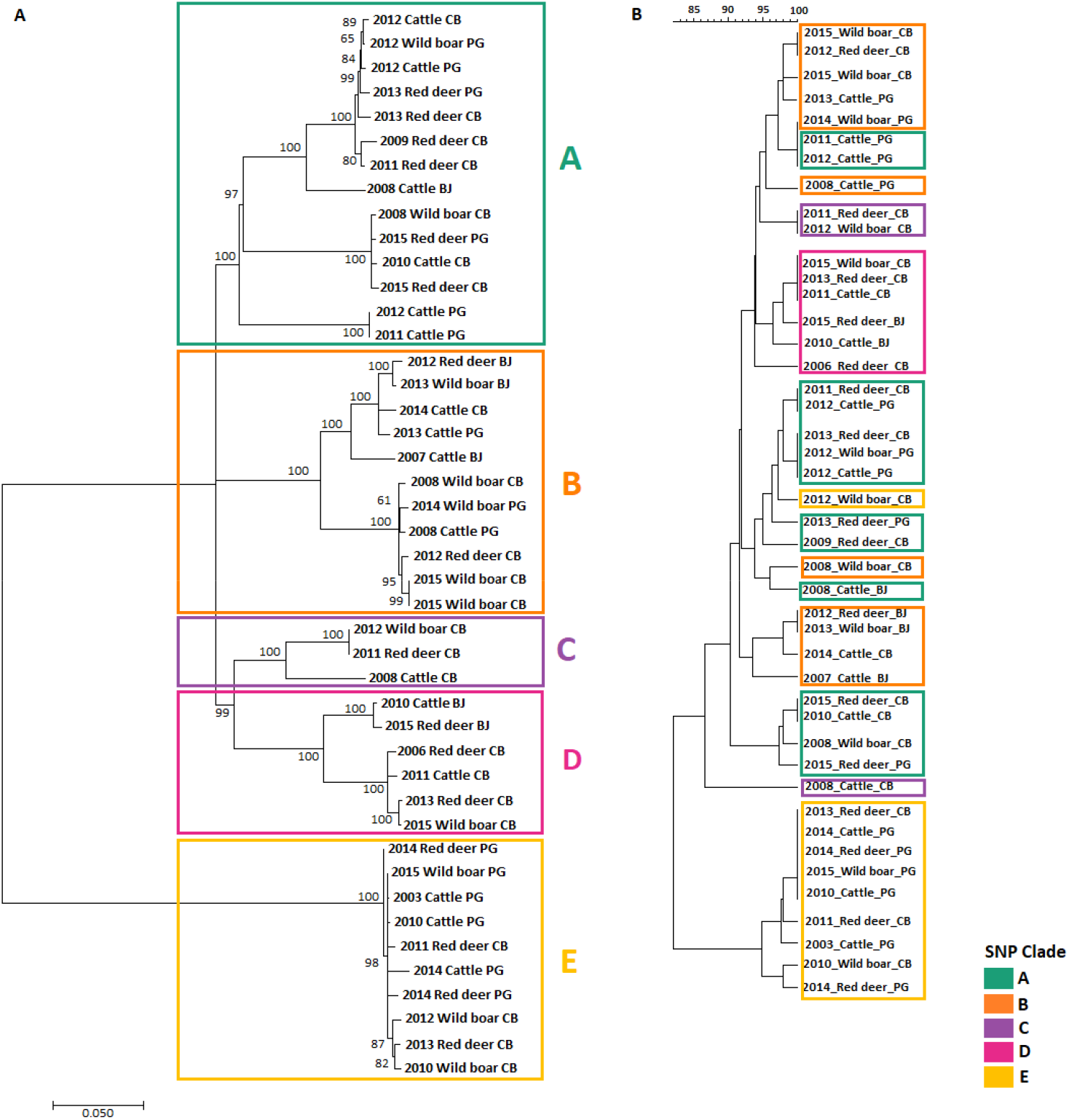
Phylogeny of *M. bovis* field strains from TB hotspots in Portugal. (A) Maximum Likelihood Tree using GTR model with input taken as an alignment file containing only informative and validated SNPs. The tree is drawn to scale, with branch lengths measured in the number of substitutions per site. (B) UPGMA tree, applying categorical option as a similarity coefficient, with input taken as the combined dataset based on spoligotyping and 8-*loci* MIRU-VNTR data. Colours identify the different SNP clades (A – dark cyan, B – orange, C – purple, D – pink and E – yellow).

**Table 1.** Number of *M. bovis* strains within each SNP clade (A to E) and identification of clade-defining and clade-monomorphic SNP sites.

| SNP clade | Total SNP sites | Clade-defining SNP sites<sup>(a)</sup> | Clade-monomorphic SNP sites<sup>(b)</sup> |
| --- | --- | --- | --- |
| A (*n*=14) | 622 | 108 | - |
| B (*n*=11) | 611 | 133 | - |
| C (*n*=3) | 320 | 184 | 49 |
| D (*n*=6) | 372 | 217 | 82 |
| E (*n*=10) | 431 | 360 | 352 |
| A to D<br>(*n*=34) | 1419 | 1411 | 106 |

(a) Polymorphic positions present only in the clade-members.
(b) Polymorphic positions present only in the clade-members and common to all members.

The topology of the SNP-based phylogenetic tree and the one based on classical genotyping techniques (combination of spoligotyping and MIRU-VNTR), agree in the large branch division between clades A to D and clade E, and in the clustering of clade D and E members (Fig. 1a and 1b).

The majority of *M. bovis* isolates (*n*=34, 77%, clades A to D) cluster within the highly structured clonal complex European 2 (Eu2) (Rodriguez-Campos et al., 2012) that is widely distributed in the Iberian Peninsula. All members from clades A to D possessed the *guaA* gene (G→A) synonymous mutation, which is the hallmark of this clonal complex (Rodriguez-Campos et al., 2012).

The clades in the upper phylogenetic branch (clades A to D) registered between 108 to 217 clade-defining SNP sites, while the lower phylogenetic branch (clade E) presented a total of 360 SNPs (Table 1). Moreover, for clades C, D and E it was also possible to identify clade-monomorphic SNP sites (i.e. polymorphic positions present only in clade members and common to them all): 49 SNPs in clade C, 82 in clade D and 352 in clade E (Table 1). When accounting for the total SNP sites registered per clade, intra-clade homogeneity (i.e. the proportion of monomorphic SNP sites within each clade) ranged from 0% to 82%, in decreasing order: clade E (82%), clade D (22%), clade C (15%) and clades A and B (0%), pointing clade E as the most homogeneous.

The differences between phylogenetic branches are also clearly expressed in a heatmap based on the absolute SNP differences between strains, which supports a very clear separation between members of clades A to D and clade E, with a high average diversity (mean SNP distance value of 384 SNPs) (Fig. 2). Grouping by host species revealed that the mean genetic difference within each group is similar (382 SNPs within cattle; 397 SNPs within red deer; and 398 within wild boar), while dissimilarities strike out when comparing the three geographic regions covered by the analyses (376 SNPs within Castelo Branco; 405 within Portalegre; and 244 within Beja) (data not shown).

**Figure 2.**
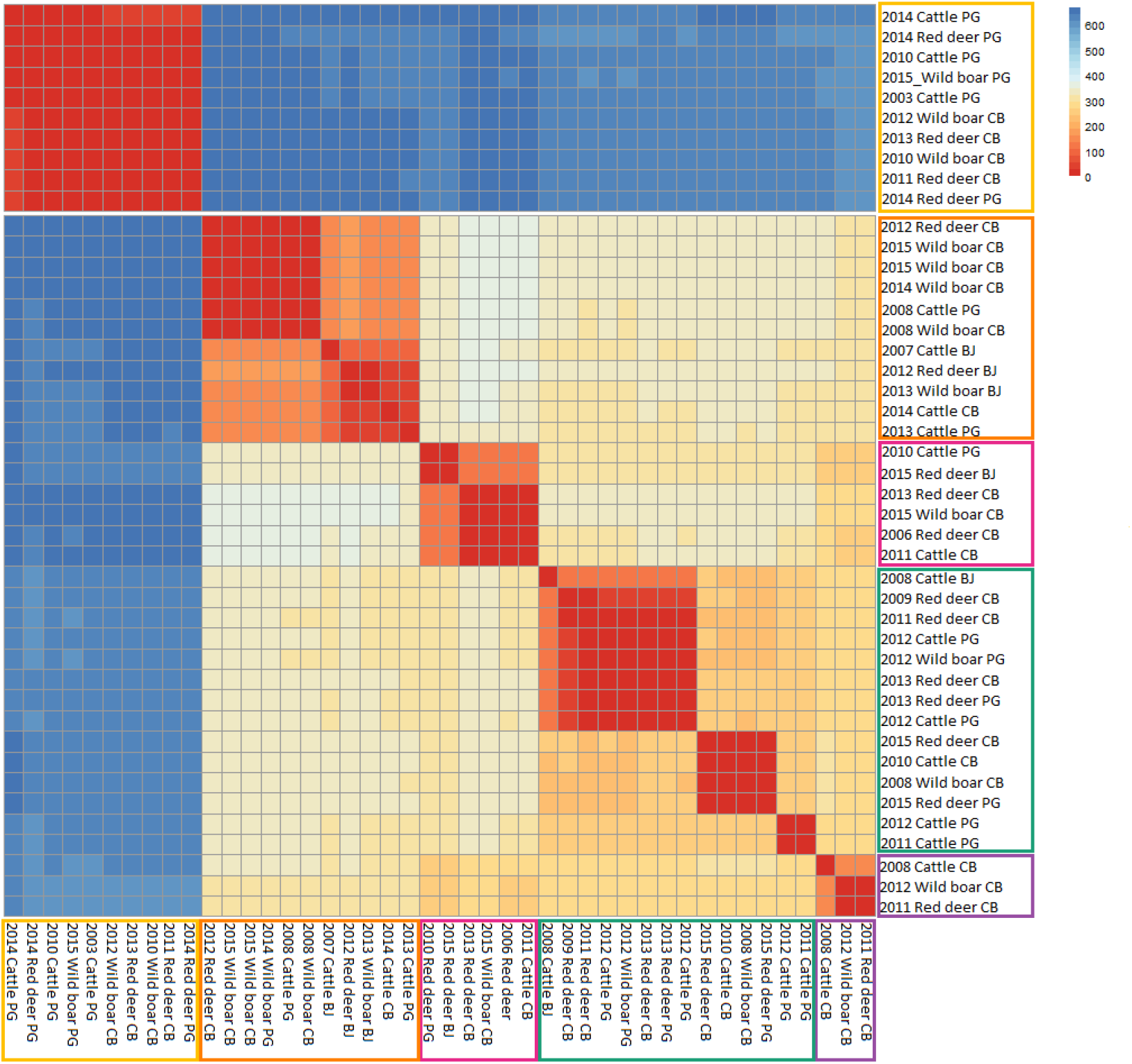
Heatmap of pairwise SNP distances based on the absolute differences of SNPs. The colours of the boxes identify the different SNP clades (A –dark cyan, B – orange, C – purple, D – pink and E – yellow).

Clades A and B were the most widely distributed, being found across the three TB hotspots under study. Clade C was exclusive of Castelo Branco, clade D was found to be absent in Portalegre, clade E was absent in Beja (Fig. 3). In contrast, similar clade distributions were found at the host species level, with strains from the three hosts clustering in the five clades (Fig. 3). When considering geographic region or host species as grouping criteria, group-defining SNP sites could also be identified for all categories under analysis, with Castelo Branco (396 SNPs) and cattle (379 SNPs) yielding the highest number of polymorphic SNPs. However, no monomorphic SNP sites were identified per host species or geographic region.

**Figure 3.**
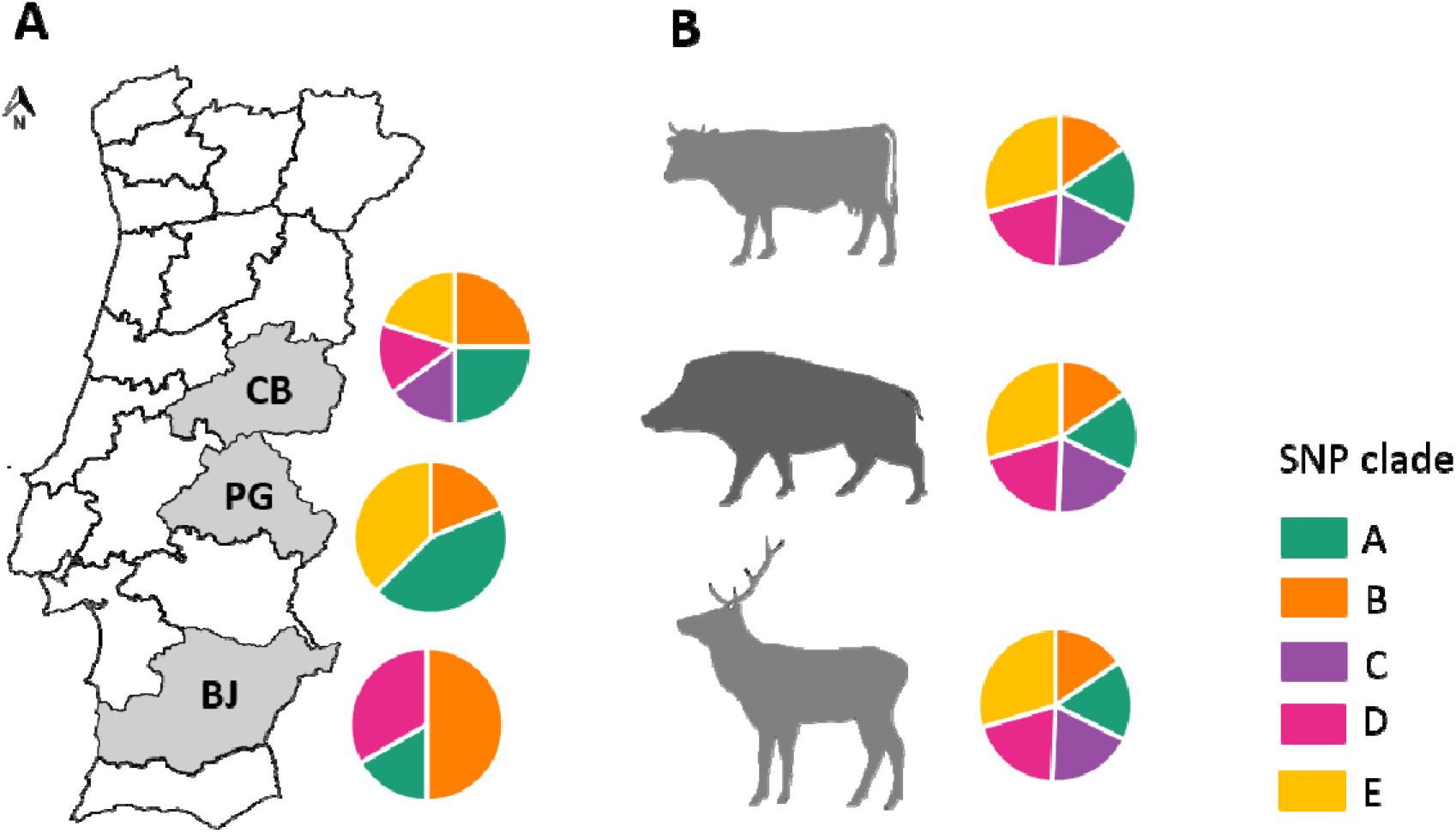
Distribution of SNP clades by geographic region (A) [Beja (BJ), Castelo Branco (CB) and Portalegre (PG)] and host species (B). The colours of the pie charts identify the different SNP clades (A –dark cyan, B – orange, C – purple, D – pink and E – yellow).

The temporal evolution of the established SNP network was assessed to get insights into the strength of relationships established through time by *M. bovis* strains, considering four progressive temporal windows and using as connection links the number of common SNPs (Fig. 4). Based on this SNP dataset, each clade could not be distinguished from the others based solely on the sampling time of the isolates (Supplementary Fig. 4). Furthermore, this analysis highlighted strong global and local networks in Portugal, with a particular focus to the strength of connections established within geographic regions (maximum shared SNPs between strains in Beja=265, Castelo Branco=370 and Portalegre=365) and between Castelo Branco and Portalegre (*n*=365) (Fig. 4).

**Figure 4.**
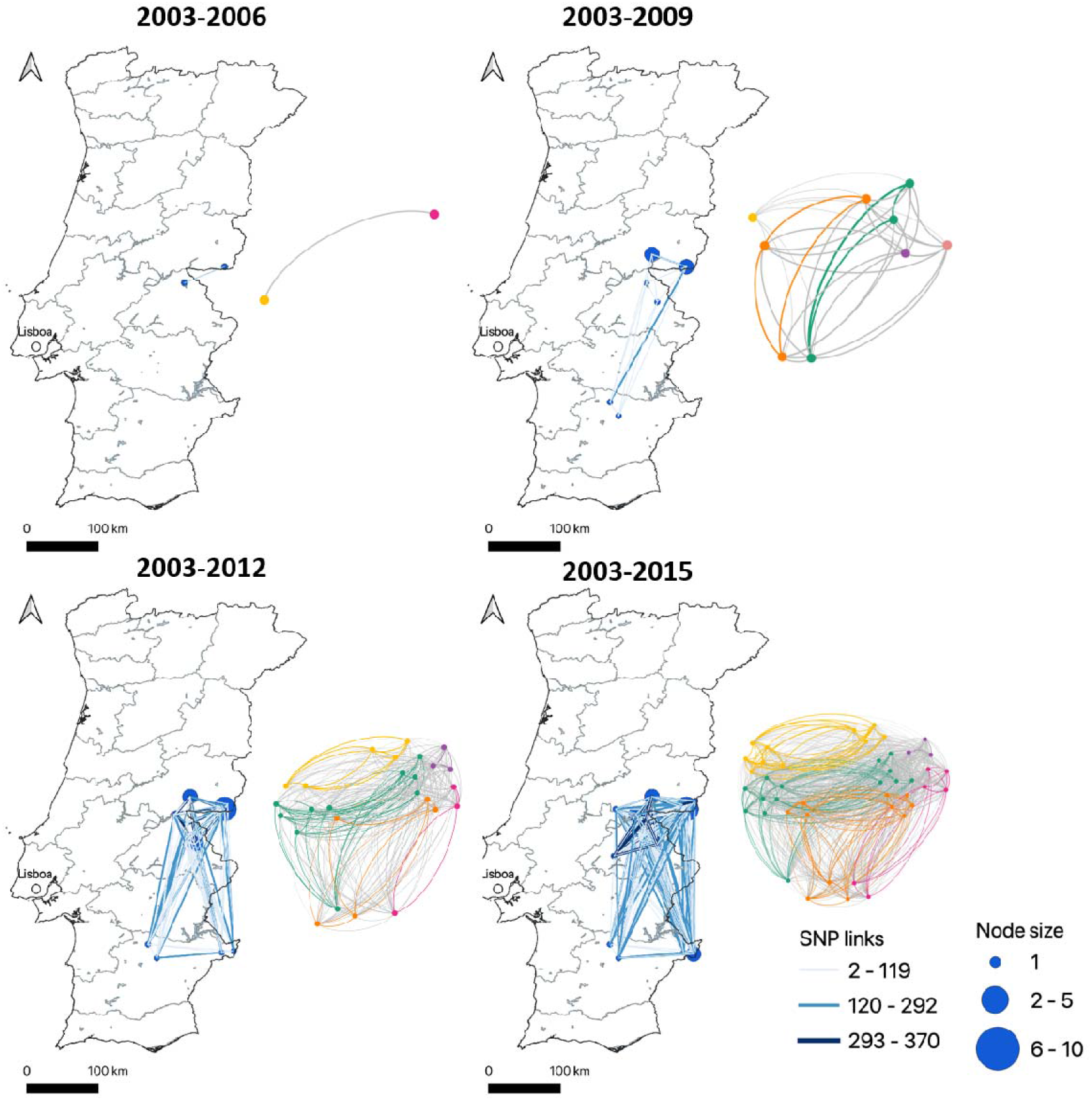
Temporal evolution of the SNP network established for the 44 *M. bovis* isolates recovered between 2003 and 2015. Four progressive time periods were considered according with the epidemiological scenario in Portugal. In the map of Portugal, the nodes represent *M. bovis* strains and the complexity of connections is based on the number of shared SNPs between strains. The side network evidences the relation established between *M. bovis* strains grouped within the same clade – each node represents one *M. bovis* strain and the colours identify the connections according with SNP clade (A – dark cyan, B – orange, C – purple, D – pink and E – yellow and grey for connections between different clades).

### 3.2. Transmission network

The global transmission network for Portuguese isolates based on the SNP alignment with 1842 positions and available metadata was constructed. The connections between *M. bovis* represent the number of SNP differences.

In the total network, 43% of individuals are linked as likely sources of infection to at least one another individual (Fig. 5). This analysis has shown that individuals could be sources of infection to up nine others, with seven (16%) linked to one host, three (7%) linked to two hosts, 22 (7%) linked to three hosts and 6 (14%) linked to four or more hosts. Links with zero SNP differences were identified within the same host species (cattle-cattle and wild boar-wild boar) and between different host species (red deer-wild boar), suggesting intra- and inter-specific transmission events (Fig. 5).

**Figure 5.**
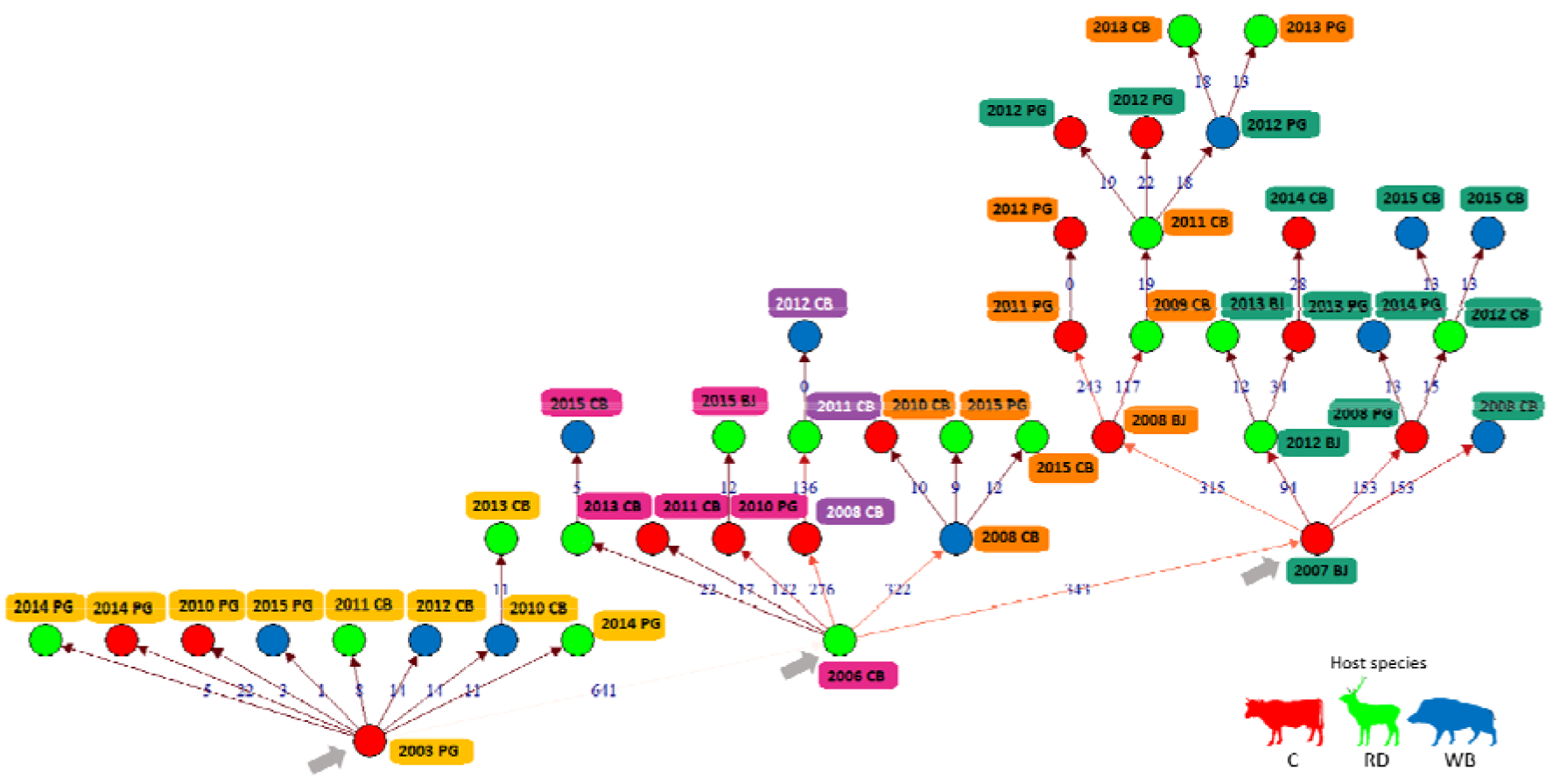
Transmission tree: reconstruction of transmission chains between cattle, red deer and wild boar, based on the SNP analyses, the year of isolation, host species and geographic region [Beja (BJ), Castelo Branco (CB) and Portalegre (PG)] of TB positive animals. The nodes are coloured by host species (cattle – red, red deer – green and wild boar – blue) and the established connections are based in SNP differences between *M. bovis* strains. Grey arrow point to the beginning of the three major branches.

Three large branches could be defined in the transmission network, with strains recovered from each one of the three TB hotspots being placed at the bottom of each branch, respectively (Fig. 5). The left subdivision of the network was exclusively composed by clade E members, while the right subdivision encompasses clades A and B affiliates. The middle branch is more diverse, with members from clades B, C and D (Fig. 5).

The number of links established by each strain was used to infer the number of secondary infections, and therefore infectivity. A stratified approach performed by host species and geographic region was represented in comparative violin plots (Fig. 6a and 6b). The graphics reveal the dispersion of the number of secondary infections, from minimum values of zero cases to a maximum of nine cases for cattle and Portalegre, if considering host species or geographic region, respectively. Plus, the higher density areas indicate that, in the majority of cases, the total number of secondary infections generated is maintained between 0 to 3, independently of host species or geographic region, with the exception of Beja (Fig. 6a e 6b). Moreover, it is also possible to conclude that cattle and Beja yield the higher median values of secondary infection (Fig. 6a and 6b). No statistically significant differences were verified when comparing secondary infections median values by host species or geographic location.

**Figure 6.**
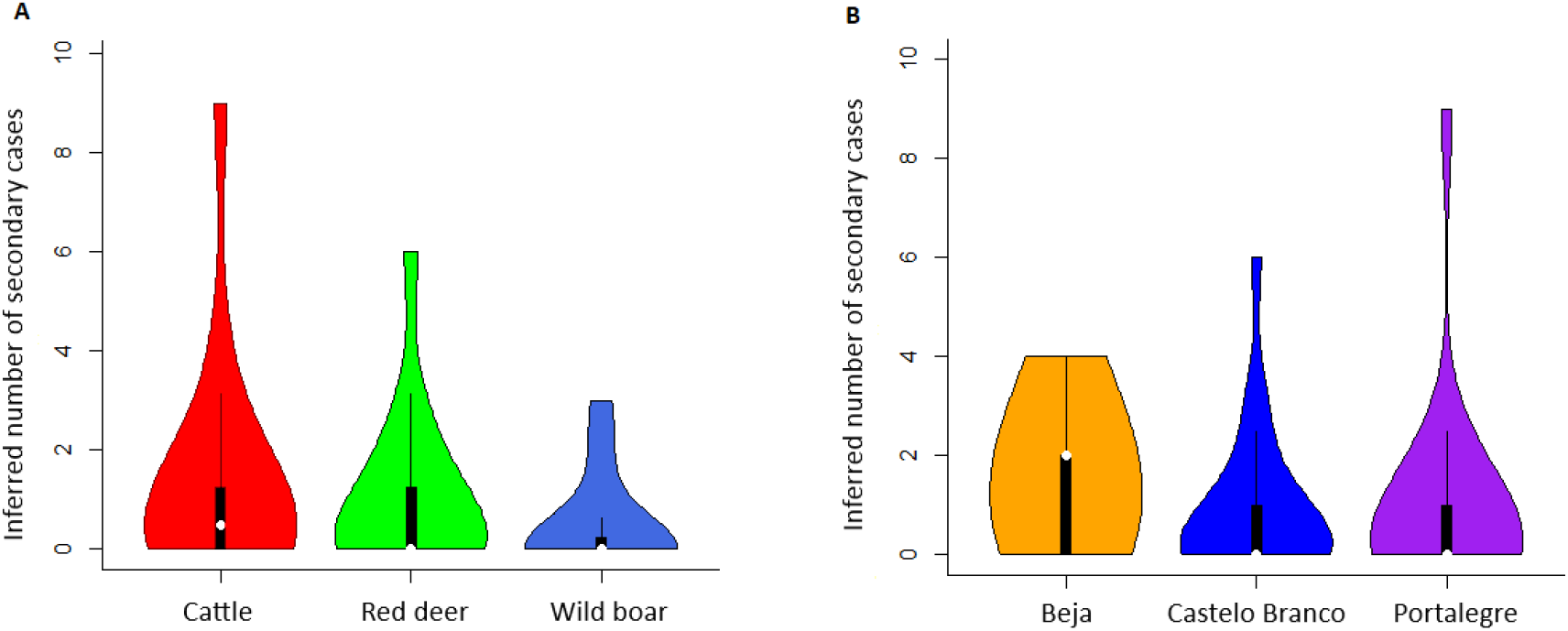
Violin plots of inferred number of secondary cases grouped by host species (A) and geographic region (B).

A comparison of pairwise genetic distance (SNPs) and geographic distance was performed, only including cases with accurate geographic coordinates (*n*=36). The Mantel test revealed no statistically significant association (simulated *p*-value= 0.541).

#### 3.2.1 Local transmission networks

Two hotspot areas in Castelo Branco (*n*=16) and Portalegre (*n*=12) were analysed and a cut-off value of 3 SNP differences in the same temporal context was used to define recent transmission events. Clustering by SNP clade is evident in both networks, with recent transmission events only involving *M. bovis* included in the same SNP clade (Fig. 7a and 7b). In Castelo Branco network, two recent transmission events were identified, with one case including individuals from different host species. In both events the wildlife hosts belong to different officially defined hunting area (Fig. 7a). Considering Portalegre’s network, one event of recent transmission was identified (Fig. 7b), with the host individuals being sampled from different herds.

**Figure 7.**
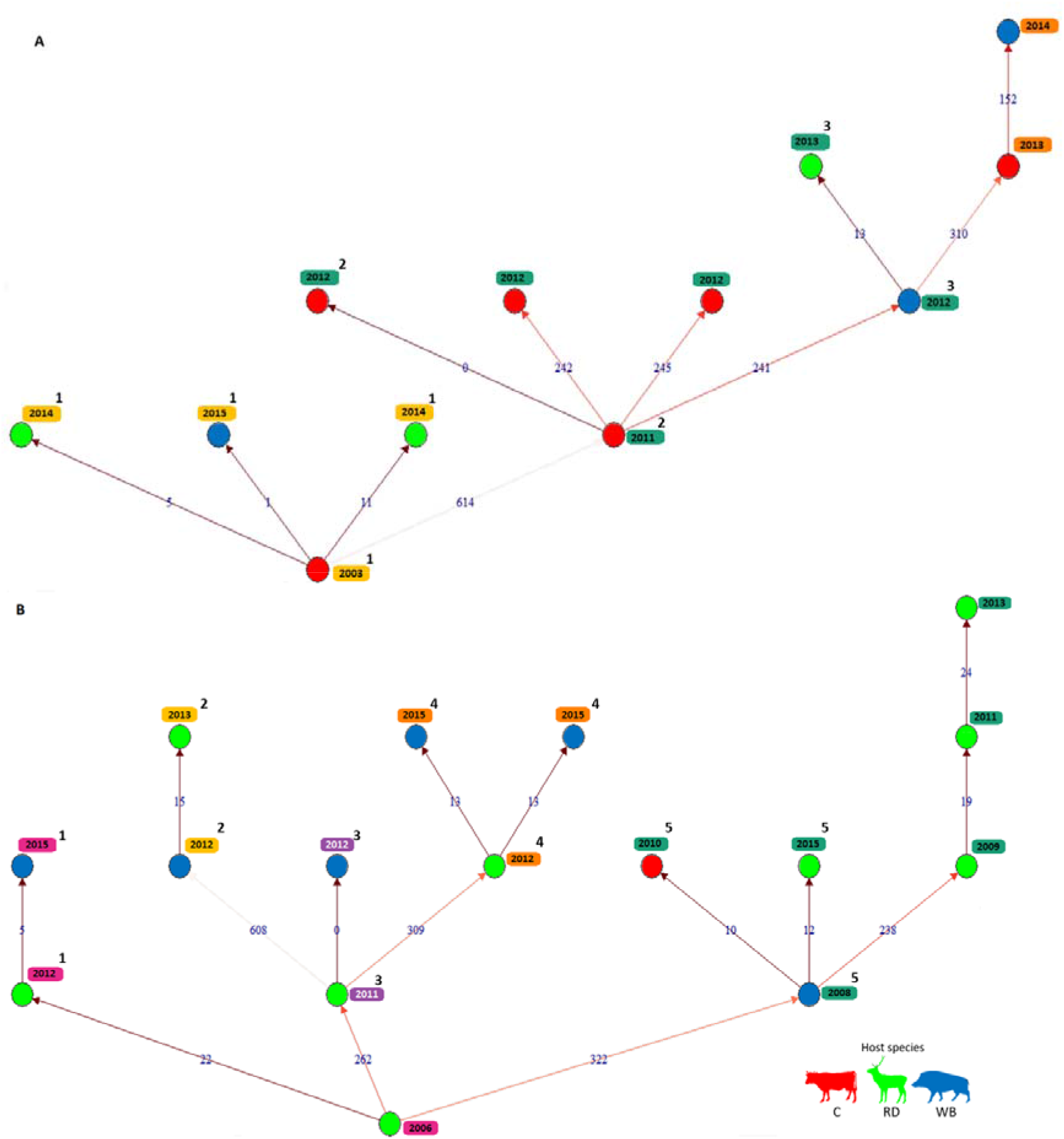
Transmission tree: reconstruction of transmission chains between cattle, red deer and wild boar, based on the SNP analyses, the year of isolation and host species in Castelo Branco (A) and Portalegre (B) of TB positive animals. The nodes are coloured by host species (cattle – red, red deer – green and wild boar – blue) and the established connections are based in SNP differences between *M. bovis*. The apostrophe numbers identify the recent transmission events (in transmission tree A, the event number “2” identifies a zero SNP link).

The Mantel test applied to Castelo Branco and Portalegre datasets revealed that there was no statistically significant association between genetic and spatial distance (simulated *p*-value=0.619 and 0.190, respectively), in agreement with findings for the global dataset from Portugal.

### 3.3. Molecular clock analysis

Before performing an evolutionary analysis, it is crucial to evaluate the clock-like structure of the entire dataset, i.e. the relationship between observed genetic distances and time. Therefore, root-to-tip regression with Tempest and DRT analysis were applied. Under a perfect clock-like behaviour, the distance between the root of the phylogenetic tree and the tips is a linear function of the tip’s sampling year, being R^2^ the degree of clock-like structure. This *M. bovis* dataset exhibits a positive correlation between genetic divergence and sampling time, however the R^2^ value obtained was low (R^2^ =0.15), suggesting that this dataset lacks a strong clock-like behaviour. Since the approach with Tempest can only be used as an exploratory analysis of temporal structure, a DRT was performed. The year of sampling was reshuffled 20 times and the clock rate of observed and randomized datasets was estimated using LSD. The clock rate estimate obtained from the observed data was 1.30× 10^-4^ [95% CI (1.0×10^-10^-3.98×10^-4^)] substitutions per-site-per-year, a value that overlaps the range of estimates obtained from the randomized sets, reinforcing that the observed data does not have a strong temporal signal (Rieux & Balloux, 2016).Therefore, considering the weak clock-like structure of this dataset, Bayesian molecular clock and evolutionary analyses were not performed.

## 4. Discussion

This work is the first study that applied WGS approaches to *M. bovis* from Portugal. The aim was to explore the fine-scale genomic signatures of field isolates and to highlight the value of phylogenetic inference and transmission networks approaches to understand the history of this pathogen at the livestock-wildlife interface.

In the current research, the SNP-based phylogeny established five main clades, with clades C, D and E presenting lower levels of intra-clade diversity and clade-specific monomorphic SNP sites that can be explored as phylogenetic markers in future diagnostic and epidemiological studies. The topology of the SNP-based phylogenetic tree agrees with the dendrogram obtained by combining classical genotyping methods, being evident that the partition between clades A-D and clade E remains well-structured and that the members of clades D and E remain together. Similar findings have been reported by other works that register clustering agreement between spoligotyping and WGS (Hauer et al., 2019; Lasserre et al., 2018). For five strains, the spoligotyping profile obtained by the reverse-hybridization method in the wet lab is not the same as the profile determined *in silico* through the vSNP pipeline, a finding that has also been reported by others (Hijikata et al., 2017; Maeda et al., 2020). This mismatch might contribute to discrepancies in phylogenetic trees topology generated with classical genotyping methods and the SNP data.

The SNP-based phylogeny revealed geographic clustering to some extent, with clade C confined to Castelo Branco and clade E being only present in Castelo Branco and Portalegre. These results are in agreement with previous work performed on this *M. bovis* population from the same geographical settings, that was based on the molecular characterization of isolates by classical genotyping methods (Reis et al., 2020). This previous study suggested the existence of five *M. bovis* ancestral populations with geographic specificities. Furthermore, a closer analysis into the temporal evolution of the SNP network evidenced strong local dynamics within *M. bovis* strains from the same geographic region, highlighting higher mean values of SNPs in common comparing with the links established between *M. bovis* strains from different geographic regions. As expected, the maximum values of common shared SNPs occurred between *M. bovis* from the bordering regions of Castelo Branco and Portalegre.

The global transmission network across regions in Portugal supports the information provided by the phylogenetic tree, with *M. bovis* isolates grouping in the same SNP clade contributing to the same transmission chains. The latency periods and long generation times of *M. bovis* can contribute to the non-agreement between phylogeny and transmission trees. Moreover, works already performed mainly with *M. tuberculosis* pointed to the variability in the number of SNPs shared between strains recovered from individuals with epidemiological links with the differential number of SNPs being used as cut-off to consider a transmission event (J. A. Guerra-Assunção et al., 2015). In the global and local transmission trees concerning this *M. bovis* population from Portugal, links with SNP differences inferior to the established cut-off value of three were identified, from the same or different host species circulating in the same spatiotemporal context, therefore suggesting recent chains of transmission. The Castelo Branco and Portalegre networks allowed a fine-scale detail of these events with the identification of three situations where the individuals were originated from different officially delimited hunting areas and herds, shedding light on the importance of animal mobility to the spread and transmission of *M. bovis* in this multi-host system. The TB eradication program implemented in Portugal is based on active and passive surveillance, but exclusively applied to the cattle population.

In contrast, surveillance in wildlife is exclusively passive and dependent upon irregular sanitary evaluation of hunter-harvested animals, meaning that *M. bovis* infection and underlying transmission might be established for long periods of time before sample collection and molecular characterization of isolates occurs, therefore limiting transmission reconstruction inferences.

In the global transmission tree, there are three large transmission chains with origin in three strains recovered from each one of the three TB hotspots, giving support to geographic clustering; plus, different degrees of inferred infectivity could be associated to each region. Beja is the most homogeneous one, presenting a lower mean value of SNP differences (244 SNPs) and a regular inferred infectivity plot; while Castelo Branco and Portalegre presented more diverse *M. bovis* populations, with higher mean SNP differences between strains (*n*= 376 for Castelo Branco and *n*= 405 for Portalegre) and with the inferred infectivity plots revealing high maximum values (six secondary cases for Castelo Branco and nine for Portalegre).

Inferences from a previous molecular approach based on two types of genomic regions - direct repeats and tandem repeats - (Reis et al., 2020) pointed the *M. bovis* population from Beja as the most ancestral, when comparing with Castelo Branco and Portalegre and populations from the latter as expanding populations. The high values of secondary cases reported in Castelo Branco and Portalegre, although in a smaller proportion, indicate higher dissemination of the pathogen. Despite the fact that this analysis only reflects the reality of these 44 cases, it mirrors the differences between expanding and stable populations.

The lack of temporal signal indicated poor correlation between genetic divergence and sampling time, stopping us to advance into evolutionary analyses. This limitation is common given the very slow mutation rate of *M. bovis* and a larger and over-sampled isolate dataset would be required to make any firm inferences regarding evolution. The large genetic distance among some isolates in the global and local transmission trees more likely reflect frequent introductions of new, more distantly related lineages into the same geographic regions.

When considering host species, the transmission network registered recent transmission events (including zero SNP differences) involving the same wildlife species or two wildlife host species, suggesting the occurrence of intra- and inter-specific transmission events. The results from inferred infectivity by host support the importance of both wildlife species in *M. bovis* transmission, with the red deer dataset presenting a higher maximum value of inferred secondary cases than wild boar, suggesting that red deer could exert a prominent role in this multi-host system. This finding is in line with previous observational studies in Portugal (Vieira-Pinto et al., 2011) and with the characteristics of the pathophysiology of TB in cervids that develop more open lesions and thus are more propense to excrete the bacilli (Cunha et al., 2012). Consequently, focus on these two wild hosts should be considered when designing new interventions aiming to improve control of animal TB.

## 5. Concluding remarks

There is clearly a need to characterise *M. bovis* lineages using clade-defining SNPs. Although limited to a modest dataset, this work confirms and reinforces the value of WGS application to the study of *M. bovis* transmission and persistence at the livestock-wildlife interface. The combination of WGS data and epidemiological information provided insights into *M. bovis* demographic history in a multi-host system, enabling the analyses of recent transmission events in several situations.

The knowledge of disease status and routes of transmission in wildlife are crucial to design and implement effective control measures. The findings reported in this work contribute to support the idea that eradication actions in the wildlife population are increasingly necessary, particularly when one considers infectivity by host species. Altogether, we provide quantitative evidence that future control measures in livestock production systems must not ignore wildlife related parameters, such as abundance, behaviour and interaction with livestock, with the possibility for a differential approach regarding red deer and wild boar. Furthermore, our fine scale molecular analyses suggest that Castelo Branco and Portalegre are to be considered priority areas of research and intervention and that adjacent livestock populations are to be tested more frequently.

Future WGS studies with a larger dataset from Castelo Branco and Portalegre, and from a broader time period, could help resolve the transmission networks and potentially provide stronger temporal signal, enabling evolutionary analyses, with estimation of evolutionary parameters, such as substitution rates, the probability of host species transition, time to the most recent common ancestor and timescales for clade divergence. Whole genome sequencing of *M. bovis* from across the world is utterly needed, not only to enlighten epidemiological scenarios, but also to build experience and tools to deal with the characteristic lack of temporal signal when slow evolving, latent microorganisms are involved. Most algorithms and simulation tools were originally developed for the evolutionary analyses of fast-evolving microorganisms (e.g. RNA viruses) and often are inadequate to deal with molecular data from highly clonal, monomorphic organisms. Future studies with a larger dataset could give further support to the use of SNP monomorphic sites as phylogenetic markers in settings where WGS may not be easy to be implemented and also contribute to understand pathogen adaptation processes to hosts.

## 6. Acknowledgements

We acknowledge the national authority for nature conservation and forests, ICNF, and the national veterinary authority, DGAV, for kindly providing metadata information. MVC personally acknowledges João Serejo for fruitful discussion on TB along the years and for outstanding field work that contributed to regular submission of wildlife TB-suspect samples. The smooth collaboration of hunting associations in field work is also recognized. This work was funded by Programa Operacional de Competitividade e Internacionalização (POCI) (FEDER component), Programa Operacional Regional de Lisboa, and Fundação para a Ciência e a Tecnologia (FCT), Portugal, in the scope of project “Colossus: Control Of tubercuLOsiS at the wildlife/livestock interface uSing innovative natUre-based Solutions” (ref. POCI-01-0145-FEDER-029783), and strategic funding to cE3c and BioISI Research Units (UIDB/00329/2020 and UIDB/04046/2020]. ACR was supported by FCT through a doctoral grant in the framework of the Applied and Environmental Microbiology Doctoral Program (PD/BD/128031/2016).

## 7. Author Contributions

MVC conceived the study. AB and TA provided the *M. bovis* isolates from the NRL. ACR performed the experimental work. SRA contributed with isolate sequencing and quality control of genome sequences. ACR, LS, and MVC thoroughly analysed the data. ACR and MVC wrote the manuscript. LS and RT contributed with critical discussion. All authors approved the final manuscript.

## 8. Data Availability Statement

Data sharing will be granted by the corresponding author upon reasonable request.

