## Supporting Information for "Phylogenomics Sheds Light on the population structure of *Mycobacterium bovis* from a Multi-Host Tuberculosis System"

**Supplementary Table 1.** Characteristics of *M. bovis* selected for WGS.

| ***M. bovis* ID** | **Sampling date** | **Host species** | **Geographic region** | **Spoligotype** | **Ancestral Populationª** |
| --- | --- | --- | --- | --- | --- |
| 220 | 2003 | Cattle | Portalegre | SB1174 | AM3 |
| 261 | 2006 | Red deer | Castelo Branco | SB0120 | AM5 |
| 601 | 2007 | Cattle | Beja | SB0295 | AM2 |
| 754 | 2008 | Cattle | Portalegre | SB0121 | AM1 |
| 769 | 2008 | Cattle | Beja | SB0119 | AM1 |
| 783 | 2008 | Wild boar | Castelo Branco | SB0121 | AM2 |
| 865 | 2008 | Cattle | Castelo Branco | SB1090 | AM4 |
| 891 | 2009 | Red deer | Castelo Branco | SB1264 | AM5 |
| 893 | 2008 | Wild boar | Castelo Branco | SB0119 | AM2 |
| 1317 | 2010 | Cattle | Beja | SB0265 | AM5 |
| 1339 | 2010 | Cattle | Castelo Branco | SB0122 | AM2 |
| 1458 | 2010 | Wild boar | Castelo Branco | SB1232 | AM3 |
| 1480 | 2010 | Cattle | Portalegre | SB1174 | AM3 |
| 1654 | 2011 | Cattle | Portalegre | SB0121 | AM1 |
| 1670 | 2011 | Red deer | Castelo Branco | SB1174 | AM3 |
| 1711 | 2011 | Red deer | Castelo Branco | SB1264 | AM1 |
| 1712 | 2011 | Red deer | Castelo Branco | SB1195 | AM5 |
| 1714 | 2011 | Cattle | Castelo Branco | SB0265 | AM5 |
| 1744 | 2012 | Wild boar | Castelo Branco | SB1264 | AM1 |
| 1746 | 2012 | Red deer | Castelo Branco | SB0121 | AM5 |
| 1758 | 2012 | Cattle | Castelo Branco | SB1264 | AM1 |
| 1769 | 2012 | Wild boar | Portalegre | SB1195 | AM5 |
| 1785 | 2012 | Red deer | Castelo Branco | SB1190 | AM5 |
| 1789 | 2012 | Cattle | Beja | SB1264 | AM1 |
| 1841 | 2012 | Cattle | Portalegre | SB0121 | AM1 |
| 1870 | 2012 | Wild boar | Portalegre | SB1264 | AM1 |
| 1915 | 2013 | Red deer | Portalegre | SB0265 | AM5 |
| 1948 | 2013 | Red deer | Castelo Branco | SB1174 | AM3 |
| 1960 | 2013 | Red deer | Castelo Branco | SB1264 | AM5 |
| 2026 | 2013 | Cattle | Castelo Branco | SB1095 | AM5 |
| 2043 | 2013 | Red deer | Portalegre | SB1264 | AM1 |
| 2067 | 2013 | Wild boar | Portalegre | SB1190 | AM5 |
| 2206 | 2014 | Cattle | Beja | SB1190 | AM1 |
| 2235 | 2014 | Red deer | Castelo Branco | SB1232 | AM3 |
| 2267 | 2014 | Cattle | Portalegre | SB1174 | AM3 |
| 2277 | 2014 | Red deer | Portalegre | SB1174 | AM3 |
| 2300 | 2014 | Wild boar | Portalegre | SB0121 | AM1 |
| 2310 | 2015 | Red deer | Portalegre | SB0122 | AM2 |
| 2313 | 2015 | Wild boar | Portalegre | SB0265 | AM5 |
| 2325 | 2015 | Red deer | Castelo Branco | SB0265 | AM5 |
| 2328 | 2015 | Red deer | Beja | SB0122 | AM2 |
| 2347 | 2015 | Wild boar | Castelo Branco | SB1174 | AM3 |
| 2395 | 2015 | Wild boar | Portalegre | SB0121 | AM5 |
| 2397 | 2015 | Wild boar | Castelo Branco | SB0121 | AM1 |

(a) Classification in ancestral population (AM1 to AM5) as described in Reis et al., 2020.

**Supplementary Table 2.** *Mycobacterium bovis* sequencing statistics details.

| ***M. bovis* ID** | **Sampling date** | **Host species^(a)^** | **Geographic region^(b)^** | **R1size** | **R2size** | **Allbam mapped reads^(c)^** | **Genome coverage^(d)^** | **Average coverage** | **Average read length** | **Unmapped reads** | **Unmapped assembled contigs** |
| --- | --- | --- | --- | --- | --- | --- | --- | --- | --- | --- | --- |
| 220 | 2003 | C | PG | 217.8MB | 230.4MB | 2394434 | 99.69 | 126.7 | 238.1 | 25077 | 2 |
| 261 | 2006 | RD | CB | 448.7MB | 466.7MB | 13725892 | 99.92 | 368.8 | 151.0 | 2146326 | 13966 |
| 601 | 2007 | C | BJ | 171.7MB | 204.8MB | 191835 | 99.73 | 98.7 | 234.2 | 22676 | 32 |
| 754 | 2008 | C | PG | 159.0MB | 171.9MB | 1359222 | 99.64 | 72.5 | 240.8 | 9193 | 4680 |
| 769 | 2008 | C | BJ | 185.4MB | 215.7MB | 2047981 | 99.68 | 108.1 | 240.2 | 19903 | 3 |
| 783 | 2008 | WB | CB | 191.2MB | 221.1MB | 2012194 | 99.66 | 105.5 | 237.8 | 22613 | 384 |
| 865 | 2008 | C | CB | 151.7MB | 162.1MB | 1410917 | 98.79 | 73.7 | 242.2 | 11861 | 366 |
| 891 | 2009 | RD | CB | 404.8MB | 419.0MB | 13142182 | 99.91 | 345.4 | 151.0 | 1898427 | 74 |
| 893 | 2008 | WB | CB | 169.3MB | 180.7MB | 188456 | 99.66 | 100.6 | 241.1 | 15639 | 4 |
| 1317 | 2010 | C | BJ | 157.2MB | 161.8MB | 1707221 | 99.59 | 91.3 | 241.5 | 13657 | 1 |
| 1339 | 2010 | C | CB | 152.4MB | 172.9MB | 1640461 | 99.63 | 87.2 | 240.5 | 13024 | 1 |
| 1458 | 2010 | WB | CB | 186.3MB | 216.5MB | 192992 | 99.67 | 103.1 | 242.2 | 19592 | 2 |
| 1480 | 2010 | C | PG | 192.4MB | 205.2MB | 205313 | 99.64 | 109.7 | 240.2 | 19865 | 2273 |
| 1654 | 2011 | C | PG | 117.7MB | 130.9MB | 615194 | 99.34 | 33.3 | 236.7 | 3968 | 60498 |
| 1670 | 2011 | RD | CB | 189.5MB | 206.5MB | 2074834 | 99.64 | 110.1 | 239.7 | 20213 | 121 |
| 1711 | 2011 | RD | CB | 161.6MB | 168.0MB | 175829 | 99.64 | 93.2 | 238.1 | 12588 | 88 |
| 1712 | 2011 | RD | CB | 121.4MB | 129.6MB | 1085013 | 99.05 | 55.6 | 241.4 | 9286 | 2030 |
| 1714 | 2011 | C | CB | 101.8MB | 110.3MB | 788996 | 99.37 | 40.5 | 241.3 | 7143 | 11013 |
| 1744 | 2012 | WB | CB | 172.5MB | 188.0MB | 1796448 | 99.69 | 92.5 | 235.1 | 14316 | 3 |
| 1746 | 2012 | RD | CB | 165.0MB | 180.6MB | 1728552 | 99.7 | 88.1 | 233 | 12827 | 1 |
| 1758 | 2012 | C | CB | 163.2MB | 175.3MB | 1732898 | 99.52 | 93.4 | 241.8 | 14928 | 1412 |
| 1769 | 2012 | WB | PG | 135.4MB | 143.5MB | 1374917 | 99.15 | 72 | 238.6 | 8851 | 1 |
| 1785 | 2012 | RD | CB | 115.5MB | 120.4MB | 940707 | 99.56 | 49.5 | 240 | 5925 | 9156 |
| 1789 | 2012 | C | BJ | 125.4MB | 133.9MB | 970309 | 99.49 | 50.8 | 234.3 | 6378 | 811 |
| 1841 | 2012 | C | PG | 168.9MB | 191.2MB | 176723 | 99.67 | 91.2 | 237.4 | 15809 | 1 |
| 1870 | 2012 | WB | PG | 130.8MB | 133.8MB | 1395146 | 99.64 | 67.7 | 223 | 12333 | 30 |
| 1915 | 2013 | RD | PG | 137.6MB | 155.4MB | 1423501 | 99.65 | 74 | 239 | 11027 | 5 |
| 1948 | 2013 | RD | CB | 153.1MB | 157.7MB | 1523134 | 99.66 | 79.4 | 237.3 | 10609 | 2 |
| 1960 | 2013 | RD | CB | 158.6MB | 177.5MB | 1712114 | 99.55 | 91.3 | 239.6 | 13976 | 1584 |
| 2026 | 2013 | C | CB | 134.4MB | 144.7MB | 1236634 | 99.61 | 64.2 | 243 | 12192 | 557 |
| 2043 | 2013 | RD | PG | 113.7MB | 121.1MB | 966319 | 99.41 | 50.3 | 241.9 | 7475 | 548 |
| 2067 | 2013 | WB | PG | 178.9MB | 193.9MB | 2004187 | 99.71 | 105.1 | 236 | 19496 | 283 |
| 2206 | 2014 | C | BJ | 162.3MB | 184.7MB | 1751017 | 99.72 | 92.7 | 238.1 | 14763 | 2505 |
| 2235 | 2014 | RD | CB | 174.0MB | 188.8MB | 1891547 | 99.65 | 100.4 | 238.9 | 16555 | 346 |
| 2267 | 2014 | C | PG | 168.4MB | 185.0MB | 890928 | 99.52 | 46.9 | 234.6 | 6772 | 4287 |
| 2277 | 2014 | RD | PG | 68.6MB | 73.6MB | 586402 | 99.36 | 29.8 | 241.5 | 4986 | 258 |
| 2300 | 2014 | WB | PG | 256.2MB | 268.0MB | 2833579 | 99.75 | 149.2 | 237.7 | 36471 | 38 |
| 2310 | 2015 | RD | PG | 132.8MB | 146.4MB | 1290912 | 99.63 | 65.8 | 234.6 | 8582 | 1136 |
| 2313 | 2015 | WB | PG | 164.8MB | 186.8MB | 1671813 | 99.65 | 85.3 | 238.3 | 17263 | 1 |
| 2325 | 2015 | RD | CB | 162.0MB | 185.5MB | 1687836 | 99.67 | 86.7 | 237.6 | 15786 | 2 |
| 2328 | 2015 | RD | BJ | 129.9MB | 150.0MB | 1220275 | 99.48 | 65.4 | 240 | 9374 | 15533 |
| 2347 | 2015 | WB | CB | 108.9MB | 124.5MB | 1135051 | 99.56 | 60.4 | 239 | 8944 | 465 |
| 2395 | 2015 | WB | PG | 216.4MB | 239.9MB | 2365469 | 99.74 | 124.4 | 237.3 | 27435 | 3 |
| 2397 | 2015 | WB | CB | 203.2MB | 215.0MB | 2171022 | 99.58 | 116.4 | 240.9 | 20636 | 1 |

(a) C - Cattle; RD - Red deer; WB - Wild boar
(b) Beja - Beja; CB- Castelo Branco; PG - Portalegre
(c) Number of successfully assembled, trimmed paired-end Illumina reads.
(d) Relative to *M. bovis* reference genome AF2122/97 (NCBI accession number NC_002945.

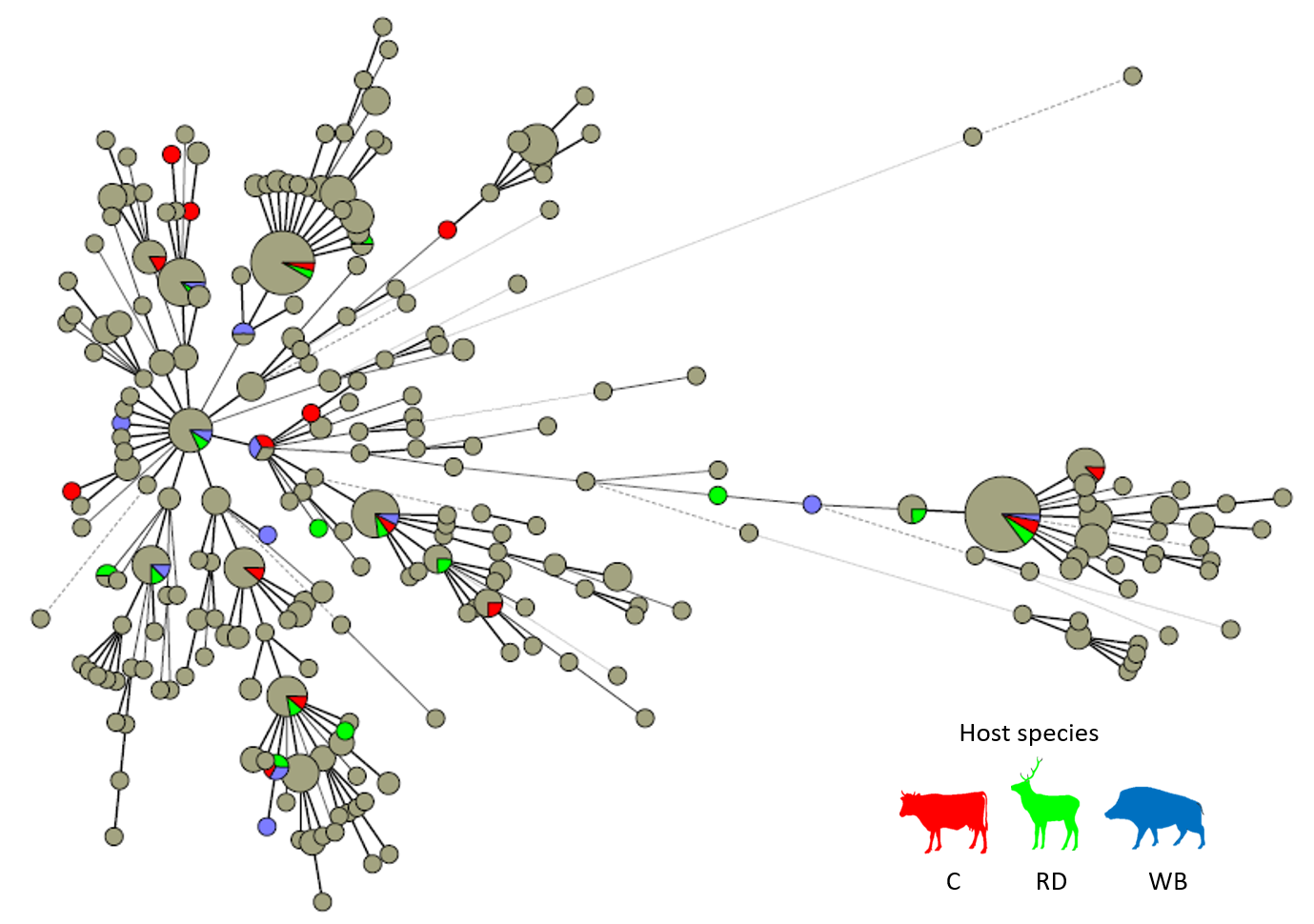
**Supplementary Figure 1.** Minimum spanning tree (MST) illustrating evolutionary relationships among *M. bovis* (*n*=487) based on spoligotyping and 8*-loci* MIRU-VNTR data, using single *locus* variant analysis. Circle size is proportional to the number of isolates within each node; and colours represent hosts species (cattle - red, red deer - green and wild boar - blue). The complexity of the lines denotes the number of differences in the spoligo-MIRU type profile between two nodes: solid lines (1, 2 or 3 differences), grey dashed lines (4 differences) and grey dotted lines (5 or more differences). Only *M. bovis* selected for WGS are coloured.

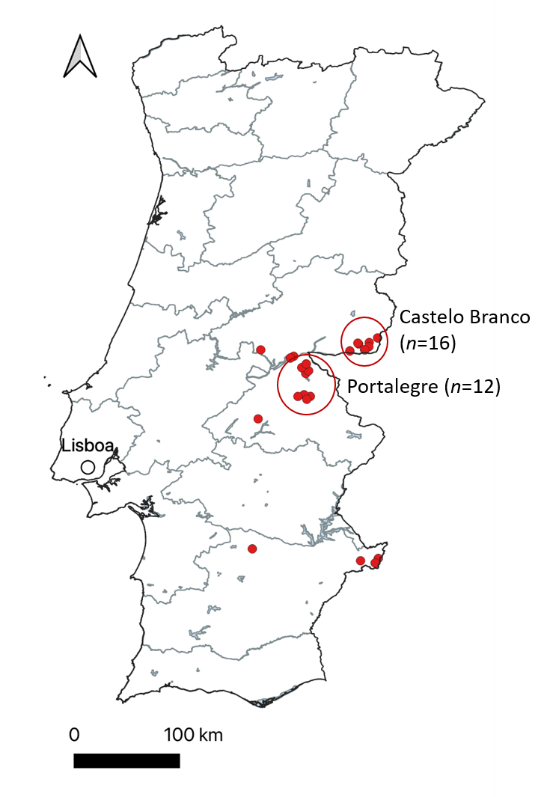

**Supplementary Figure 2.** Map of Portugal representing the *M. bovis* isolates with geographic coordinates available. The red dots represent the location of each *M. bovis.* The circles identify the two hotspot areas.

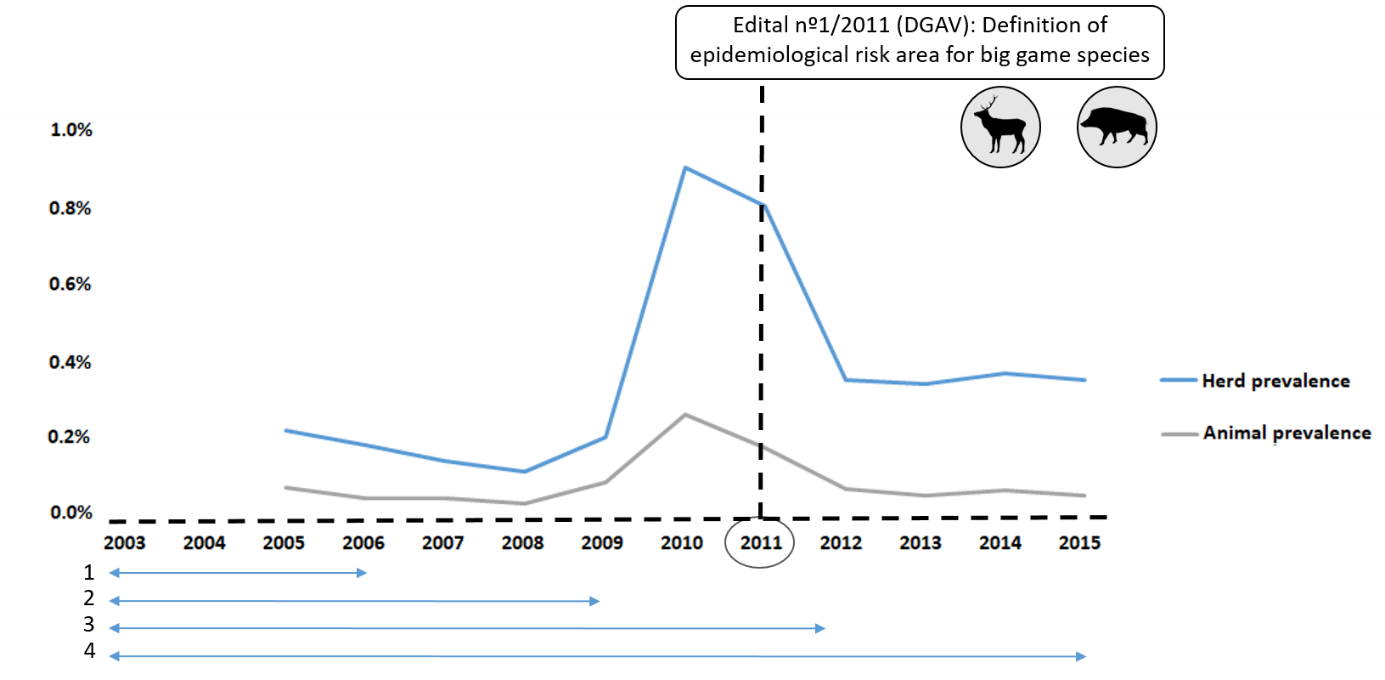

**Supplementary Figure 3.** Time line of animal tuberculosis in Portugal. The curves represent the evolution of animal TB epidemiological indicators, for cattle population, in mainland Portugal (2003-2017) [Adapted from Relatório Técnico de Sanidade Animal, DGAV (2015)]. The year 2011 marks the definition of epidemiological risk area for big game species. The four progressive time periods considered in this work are evidenced under the time line.

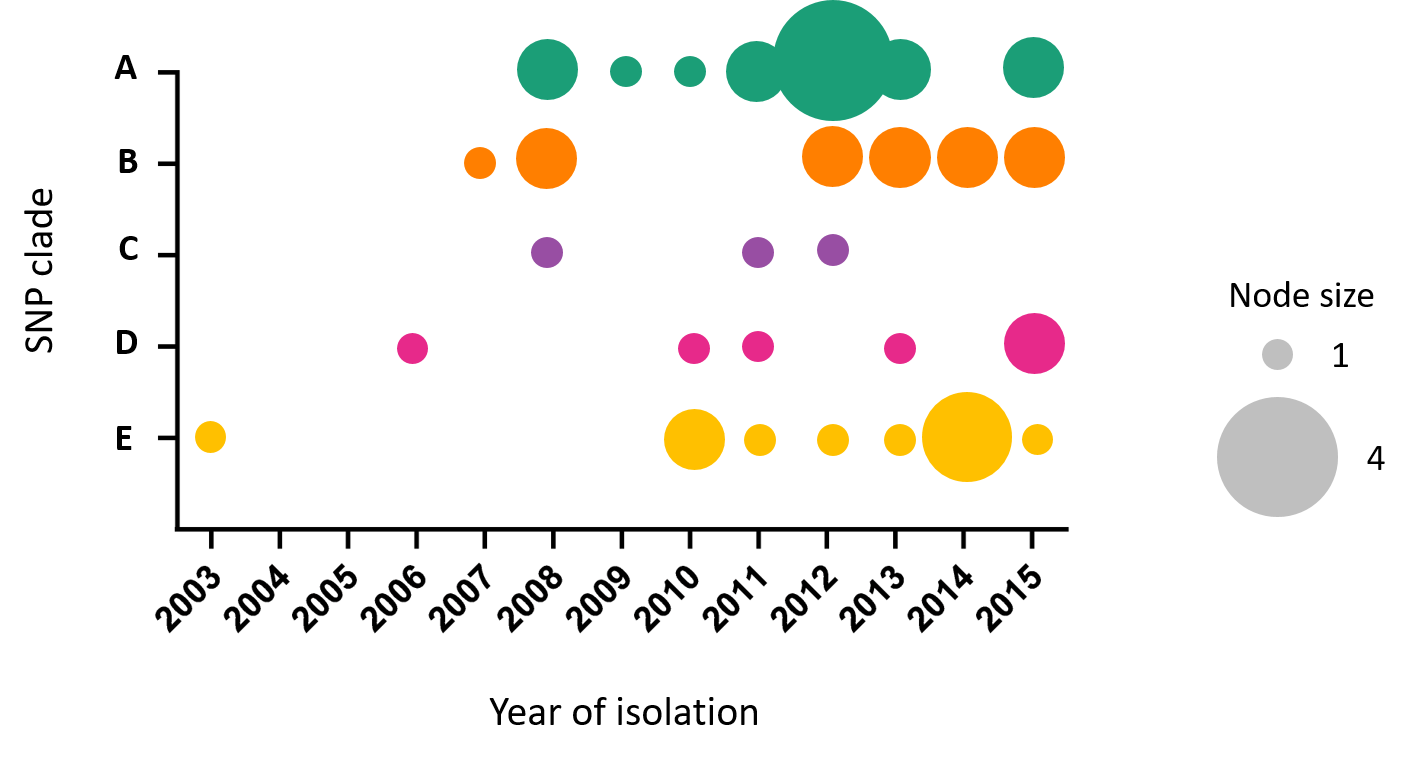

**Supplementary Figure 4.** Temporal distribution of *M. bovis* identified by SNP clade per year. Node size is proportional to the number of *M. bovis* strains within each node.
